# The CryoEM structure of human serum albumin in complex with ligands

**DOI:** 10.1101/2024.02.21.581427

**Authors:** Claudio Catalano, Kyle W. Lucier, Dennis To, Skerdi Senko, Nhi L. Tran, Ashlyn C. Farwell, Sabrina M. Silva, Phat V. Dip, Nicole Poweleit, Giovanna Scapin

**Author notes:** Rowan SOM, 1 Medical Center Dr., Stratford, NJ 08084.

## Abstract

Human serum albumin (HSA) is the most prevalent plasma protein in the human body, accounting for 60% of the total plasma protein. HSA plays a major pharmacokinetic function, serving as a facilitator in the distribution of endobiotics and xenobiotics within the organism. In this paper we report the cryoEM structures of HSA in the apo form and in complex with two ligands (salicylic acid and teniposide) at a resolution of 3.5, 3.7 and 3.4 Å, respectively. We expand upon previously published work and further demonstrate that sub-4 Å maps of ∼60 kDa proteins can be routinely obtained using a 200 kV microscope, employing standard workflows. Most importantly, these maps allowed for the identification of small molecule ligands, emphasizing the practical applicability of this methodology and providing a starting point for subsequent computational modeling and in silico optimization.

## Introduction

Human serum albumin (HSA) is the predominant protein found in blood^1^. Its synthesis originates in the hepatocyte, starting as a precursor protein known as pre-pro-albumin. This precursor undergoes modification into pro-albumin within the lumen of the endoplasmic reticulum. The maturation is facilitated by furin cleaving a 6-amino acid long oligopeptide at the N-terminal in the trans-Golgi apparatus. This releases the mature plasma protein into the intravascular space, with a typical concentration ranging from 35 to 50 g/L^1^. HSA, with its abundant presence, serves a variety of functions, including the regulation of osmotic pressure, modulation of fluids distribution between body compartments, control of redox potential, and buffering of pH levels. Additionally, it acts as a carrier protein of endobiotic and xenobiotic substances within the organism, exhibiting an extraordinary binding capacity to molecules like fatty acids, hormones, free radicals, and drugs^2–11^.

HSA is a single polypeptide chain of 585 amino acids, with a molecular weight of 65 kDa. The secondary structure of HSA is primarily helical, composed of 67% **α**-helices, loops and turns, lacking **β**-sheet elements. The first three-dimensional structure, determined by X-ray crystallography, was published in 1992 (PDB ID: 1UOR)^12^ at a resolution of 2.8 Å. Since then, over 130 crystal structures of HSA have been deposited in the PDB, and very recently one cryogenic electron microscopy (cryoEM) structure of the apo form to 3.7 Å has also been reported (PDB ID: 8Q3F)^13^. The ternary globular shape of HSA is preserved and stabilized by the existence of 35 cysteines, engaging in 17 disulfide bridges (cystines). Notably, the sole free cysteine residue at position 34 is putatively dedicated to maintaining the redox balance within the plasma. At physiological pH it has a net charge of negative 15, owing to its numerous ionizable residues, making it an effective plasma buffer. The multidomain organization of HSA is reflected in the extraordinary ligand binding properties. It can bind many drugs and can contribute to their distribution, metabolism, and elimination in the body, affecting their pharmacokinetics, bioavailability, and therapeutic efficacy^6,9,11,14^.

Understanding the binding interactions between drugs and HSA is crucial in drug discovery and development^7,14–20^. The binding affinity and binding kinetics of drugs to HSA are usually evaluated during the drug development process, to assess their potential off-target effects and optimize their pharmacokinetic properties. Structural knowledge of a drug-HSA complexes can greatly help in defining specific interactions that can be modulated or changed to alter the pharmacokinetic properties of molecules and in some cases improve safety and efficacy^21–24^.

With a molecular weight of about 65 kDA, HSA is much smaller than what is considered a size suitable for routine structure determination by cryoEM. While CryoEM structures of molecules smaller that 100 kDa in size are available^25–28^, many challenges remain in studying small proteins using cryoEM; for example, the low sample size results in a lower signal-to-noise ratio in the collected images, which makes particle picking difficult, and the lack of structural markers renders the alignment in 2D and in 3D more difficult to achieve^29–31^. On the other hand, many drug targets are relatively small proteins or smaller subdomains of larger complexes, such as GPCRs, kinases, phosphodiesterases, and proteases^32–34^ and being able to use cryoEM to determine structural information on the native proteins would be a useful addition to the current structural toolset. In response to these challenges, we sought to address two key questions. First, could we successfully resolve a cryoEM structure of a sub-70 kDa asymmetric protein using a basic 200 kV instrument? Secondly, could we reach the resolution necessary to confidently place a ligand in its binding site(s)? The outcomes of our efforts are presented in this paper, which describes the cryoEM structures of unliganded (apo) HSA and HSA in complex with salicylic acid (SAL) and teniposide to 3.5, 3.7, and 3.4 Å resolution, respectively. These results not only demonstrate the feasibility of routinely studying small proteins using cryoEM but also show the ability to identify small molecule ligands in near-atomic resolution maps. This structural information proves valuable for subsequent modeling and *in silico* improvement of compounds, underscoring the potential impact on advancing drug discovery and development processes.

## Results and discussion

### Establishment of general vitrification, data collection and data processing conditions

As for all structural techniques (including NMR, mass spectrometry and crystallography), sample quality plays a pivotal role in the success of a single particle analysis (SPA) cryoEM campaign and the availability of consistently reliable samples is essential in establishing a robust platform for routine determination of 3D structures^35–38^. For this reason, we conducted an initial quality control assessment of the purchased HSA, using size exclusion chromatography, mass photometry, and negative staining analysis. We also considered the sample’s performance after extended storage at both 4°C and −80°C under various concentrations (see SI for more details). The results indicated that the sample was always monomeric, highly homogeneous, and monodisperse (Supp Figure 1). These findings provide strong evidence that the protein could be routinely prepared and employed for the precise determination of high-resolution structures and related research efforts.

**Figure 1:**
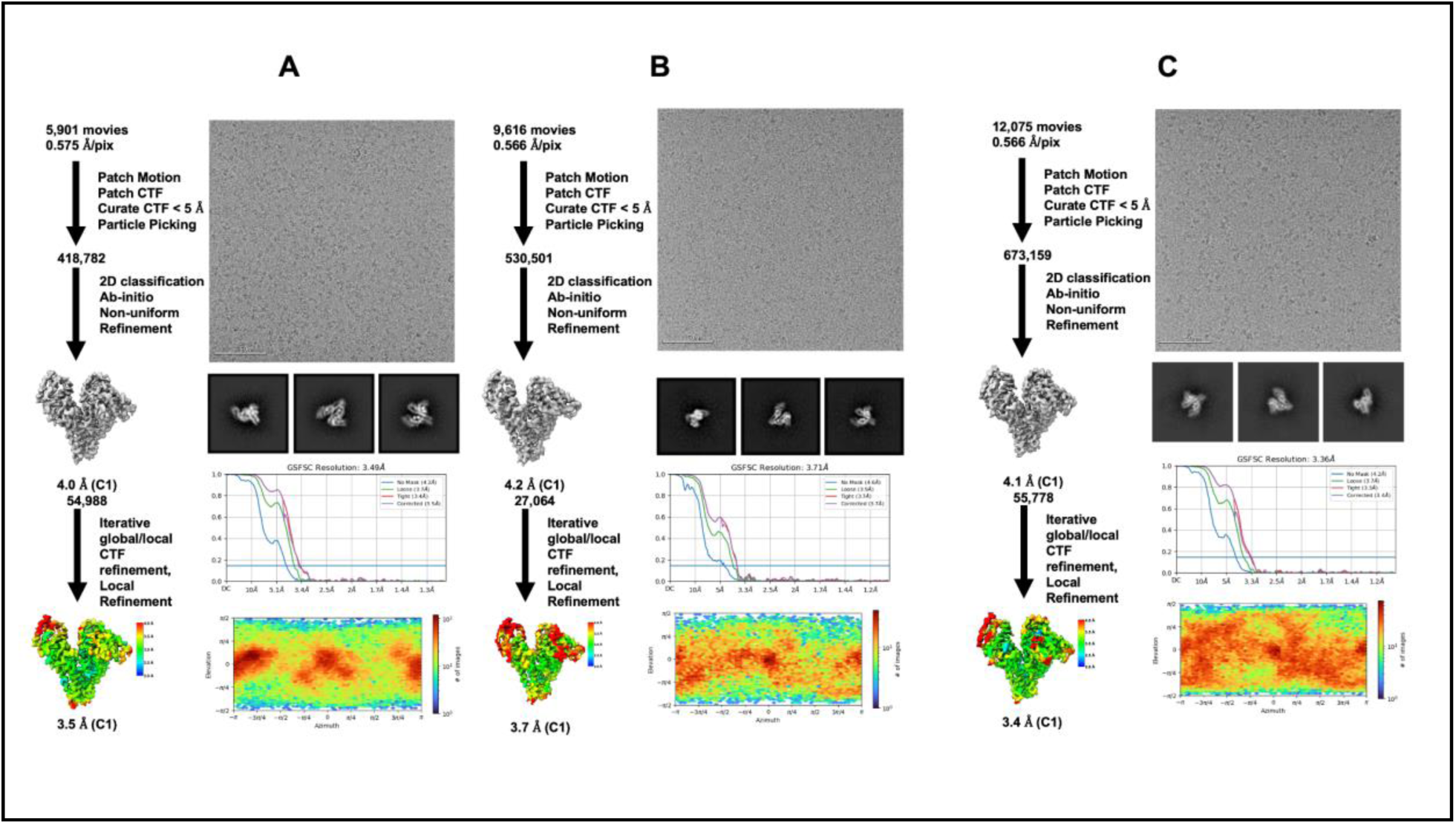
Summary of data processing. Processing workflow, exemplary micrograph, 2D classes, overall map, FSC curve and viewing distribution as calculated by cryoSPARC^51^ for **a** the apo HSA; **b** HSA bound to salicylic acid and **c** HSA bound to teniposide.

As a subsequent step, our focus shifted towards implementing a robust cryoEM 3D structure determination workflow, encompassing the entire process from vitrification to structure solution, for the unliganded sample. This step was crucial for several reasons. First, it served as an additional quality control check for the protein since the results of negative staining alone cannot reliably predict how the protein behaves during vitrification. Second, having a standardized process in place allowed us to establish a control baseline, which would be invaluable in cases of unexpected behavior in the presence of compounds or when dealing with a new batch of protein. Lastly, this procedure provided the opportunity to explore a diverse range of conditions, including variations in protein concentration, grid type, and vitrification methods, to establish the most effective parameters, while simultaneously evaluating the reproducibility of the method. These insights then served as a foundational reference for vitrifying complexes in subsequent experiments (SI and Supp Table 1).

**Table 1.** Blotting Parameters and Glacios Imaging Conditions.

| **Table 1. Blotting Parameters and Glacios Imaging Conditions** |  |  |  |
| --- | --- | --- | --- |
| Sample | HSA WT | HSA Salicylic Acid | HSA Teniposide |
| Blot Time | 10 seconds | 3 seconds | 3 seconds |
| Blot Force | 10 | 25 | 5 |
| Wait Time | 0 seconds | 0 seconds | 0 seconds |
| Imaging Mode | Nanoprobe | Nanoprobe | Nanoprobe |
| Detector Mode | Counting | Counting | Counting |
| Electron Voltage | 200 kV | 200 kV | 200 kV |
| Nominal defocus | -0.5 $\mu\text{m}$ to -1.5 $\mu\text{m}$ | -0.5 $\mu\text{m}$ to -1.5 $\mu\text{m}$ | -0.5 $\mu\text{m}$ to -1.5 $\mu\text{m}$ |
| Nominal magnification | 240000 | 240000 | 240000 |
| Pixel size | 0.575 Å | 0.566 Å | 0.566 Å |
| Total dose | 40.96 $\text{e}^-/\text{\AA}^2$ | 36.43 $\text{e}^-/\text{\AA}^2$ | 36.43 $\text{e}^-/\text{\AA}^2$ |
| Total exposure time | 3.5 seconds | 3.5 seconds | 3.5 seconds |
| EER internal frames | 840 | 840 | 840 |
| Imaging | Image shift | Image shift | Image shift |
| Total number of Images | 5901 | 9616 | 12075 |

All grids underwent screening, and data collection was carried out on suitable grids to further characterize the vitrification conditions (SI). As anticipated, ice thickness proved to be a critical factor for the success of the project^39^. The grid that yielded the 3.5 Å structure exhibited an average ice thickness of 20 nanometers, as evaluated by Appion^40^, Supp Figure 2). Another key factor was the utilization of a nominal magnification of 240K, corresponding to a pixel size of 0.56 Å during data collection^41^. These established vitrification conditions, data collection parameters, and data processing steps were subsequently applied to the liganded samples with minimal modifications.

**Figure 2:**
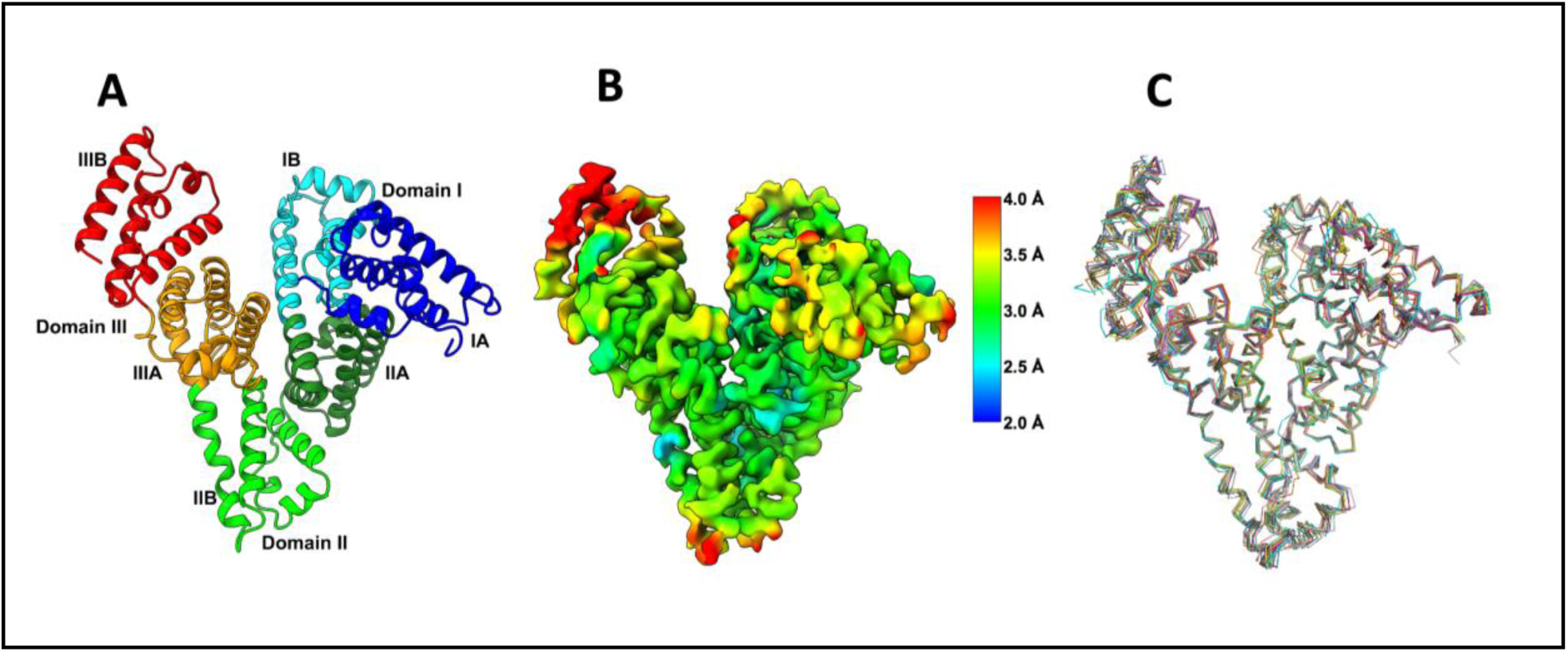
The unbounded HSA. **a** Identifications of the domains in the HSA structure. **b** Map at 3.5 Å colored according to the local resolution; **c** alignment of the X-ray structures for unbounded HSA (PDB ID: 1AO6; 1UOR; 1BM0; 7VR0; 6M5D; 7FFR; 7FFS; 6M4R; 7DJN; 1E78; 5Z0B; 4G03; 4G04; 4K2C) only the ca-traces are displayed; the alignment was carried out in Pymol^60^.

### CryoEM structure of apo HSA

The final map for HSA was reconstructed to a nominal resolution of 3.5 Å (Fig 1, Table 1 and Table 2). The overall structure can be described as an asymmetric heart-shaped molecule with a diameter of approximately 7 nm (80 x 80 x 30 Å) containing three homologous **α**-helical domains usually indicated as I (1-195), II (196-383), and III (384-585) (Fig R1). The three domains are structurally similar, each having ten antiparallel helices subdivided into two subdomains, A and B. Each subdomain A is constituted of six helices (h1-h6) and each subdomain B of four helices (h7-h10). HSA has multiple binding sites (up to 8)^5,8^, including well-characterized drug-binding sites located in domain IIA and IIIA called Sudlow’s Sites (Figure 2a)^42^. The local resolution of the cryoEM map (Fig 2B) shows that while the overall resolution is 3.5 Å, many regions of the map, including the two Sudlow’s sites, exhibit a local resolution of 2.5-3.2 Å, with well-defined side chains (Supp figure 3). The quality of the map facilitated the straightforward building of the HSA model into the available density, and subsequent refinement (refinement statistics are reported in Table 2). The resulting model is basically identical to the reported crystal structures, having an overall RMSD on CA of ∼0.8 Å with respect to structure 1AO6 (which will be used for comparison throughout). The major differences between the cryoEM structure and the x-ray structure are located in domain IA and IIIB. While these differences can be attributed to the presence of crystal contacts (for example domain IIIA in 1AO6 is in close contact with domain 1B of the second molecule present in the asymmetric unit), they may reflect intrinsic biological properties of HSA: a quite large amount of local flexibility is indeed discernible in domains IA, IIA and IIIB when overlaying the 15 crystal structures of unbound HSA downloaded from the PDB (depicted in Figure 2C). Notably, the local resolution map calculated from the cryoEM data (Figure 2B) also allows for the visualization of the different conformational flexibility within the various domains, with domain IIIB showing the highest flexibility, followed by domains IA and IIB. A single cryoEM map, even at medium resolution, can thus provide valuable information about dynamics and flexibility without the need of an atomic model. Such information can be useful in understanding biological data.

**Figure 3:**
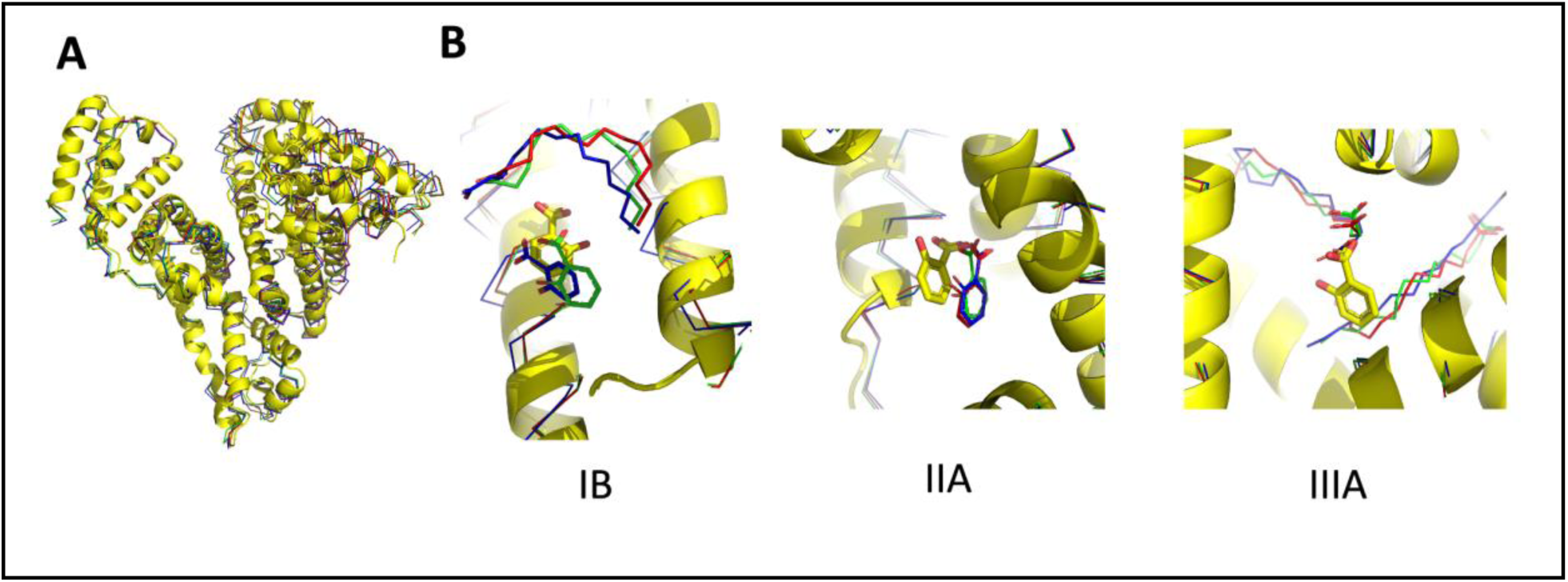
CryoEM structure of salicylic acid bound to HSA. Comparison between the crystal structures (PDB ID: 2I2Z, red; 2I30, green; 3B9M, blue) and the cryoEM structure (yellow) of the complexes between HSA and salicylic acid. **a** Overlay of the ca-traces for the four structures: the major differences are in domain I and II, reflecting the presence of bound myristate in the X-ray structures. **b** The individual binding sites for SAL; in the crystal structures fatty acids are present in both sites IB and IIIA (as well close to site IIA), and the overall shape of site IIA in the crystal structures and the cryoEM structure is different because of the conformational rearrangement induced by the fatty acid.

**Table 2:** Refinement Statistics.

| Table 2: Refinement Statistics |  |  |  |
| --- | --- | --- | --- |
|  | HSA WT | HSA Salicylic Acid | HSA Teniposide |
| Total extracted picks | 418,782 | 530,501 | 673,159 |
| Final Particles (no.) | 54,988 | 27,064 | 55,778 |
| Symmetry | C1 | C1 | C1 |
| FSC 0.143 | 3.5 Å | 3.7 Å | 3.4 Å |
| Model |  |  |  |
| Chains | 1 | 2 | 2 |
| Atoms | 4623 (Hydrogens: 0) | 4653 (Hydrogens: 0) | 4669 (Hydrogens: 0) |
| Residues | Protein: 581 | Protein: 581 | Protein: 581 |
| Water | 0 | 0 | 0 |
| Ligands | 0 | SAL:3 | 9TP:1 |
| Bonds (RMSD) |  |  |  |
| Length (Å) (# > 4sigma) | 0.003 (0) | 0.003 (0) | 0.003 (0) |
| Angles (°) (# > 4sigma) | 0.587 (1) | 0.662 (4) | 0.664 (1) |
| MolProbity score | 1.69 | 1.63 | 1.94 |
| Clash score | 6.11 | 8.69 | 10.82 |
| Ramachandran plot (%) |  |  |  |
| Outliers | 0.00 | 0.00 | 0.00 |
| Allowed | 5.18 | 2.94 | 5.70 |
| Favored | 94.82 | 97.06 | 94.30 |
| Ramachandran plot Z-score (RMSD) |  |  |  |
| Whole (N=579) | 1.47 (0.36) | 1.75 (0.35) | 1.17 (0.36) |
| Helix (N=413) | 1.75 (0.26) | 1.68 (0.26) | 1.21 (0.26) |
| Sheet (N=0) | --- (---) | --- (---) | --- (---) |
| Loop (N=166) | -1.01 (0.48) | -0.03 (0.50) | -0.16 (0.53) |
| Rotamer outliers (%) | 0.00 | 0.00 | 0.00 |
| Cbeta outliers (%) | 0.00 | 0.00 | 0.00 |
| Peptide plane (%) |  |  |  |
| Cis proline/general | 4.2/0.0 | 4.2/0.0 | 4.2/0.0 |
| Twisted proline/general | 0.0/0.0 | 0.0/0.0 | 0.0/0.0 |
| CaBLAM outliers (%) | 2.25 | 1.39 | 2.77 |
| ADP (B-factors) |  |  |  |
| Iso/Aniso | 4623/0 | 4653/0 | 4669/0 |
| Protein<br>(min/max/average) | 20.8/134.6/56.8 | 54.4/175.6/94.2 | 48.5/143.4/81.9 |
| Ligand<br>(min/max/average) |  | 73.4/84.0/80.0 | 88.1/88.1/88.1 |
| **Data** |  |  |  |
| Box pixel size | 440/440/440 | 440/440/440 | 440/440/440 |
| Angles (°) | 90.00, 90.00, 90.00 | 90.00, 90.00, 90.00 | 90.00, 90.00, 90.00 |
| Supplied Resolution (Å) | 3.5 | 3.7 | 3.4 |
| Resolution Estimates (Å) | Masked-Unmasked | Masked-Unmasked | Masked-Unmasked |
| d FSC (half maps; 0.143) | 3.5-3.7 | 3.8-3.9 | 3.5-3.6 |
| d 99 (full/half1/half2) | 3.7/1.2/1.2<br>3.7/1.1/1.1 | 3.9/1.1/1.1-3.9/1.1/1.1 | 3.6/1.1/1.1-3.5/1.1/1.1 |
| d model | 3.6-3.6 | 3.8-3.8 | 3.6-3.5 |
| d FSC model<br>(0/0.143/0.5) | 3.3/3.4/3.6-3.4/3.5/3.7 | 3.5/3.6/3.9-3.6/3.7/4.0 | 3.2/3.3/3.6-3.3/3.4/3.8 |
| Map min/max/mean | -0.24/0.41/0.00 | -0.18/0.33/0.01 | -0.20/0.38/0.00 |
| Model vs. Data |  |  |  |
| CC (mask) | 0.84 | 0.81 | 0.83 |
| CC (box) | 0.81 | 0.83 | 0.79 |
| CC (peaks) | 0.76 | 0.73 | 0.72 |
| CC (volume) | 0.81 | 0.79 | 0.82 |
| Mean CC for ligands | --- | 0.73 | 0.69 |

Emerging evidence points to the importance of HSA dynamics not only in the binding of multiple ligands^43^, but also for the interactions with receptors^44^. Furthermore, it has been reported that HSA has an inhibitory, chaperon-like function, regulating the formation and aggregation of amyloid β fibril in a fatty-acid and cholesterol dependent manner. The intrinsic flexibility of HSA is proposed to be advantageous in forming promiscuous interactions with various partners^45^. It would be interesting to see if cryoEM analysis of a FA saturated HSA shows the same pattern of flexible domains, or if the presence of FA stabilizes in part the protein.

### CryoEM structure of salicylic acid (SAL) bound HSA

The final map for HSA in complex with SAL was reconstructed to a nominal resolution of 3.7 Å (Figure 1, Table 1 and Table 2). The structure was determined in the absence of fatty acids, and the overall conformation as well as the relative position of the three subdomains closely resemble those of the apo protein. In contrast, the structures of HSA:SAL available in the PDB (PDB ID: 2I30; 2I2Z; 3B9M) feature myristate binding, and the presence of these lipids not only influences the overall conformation of the protein (Figure 3A), but also impacts the salicylic acid binding sites shape and dimensions (Figure 3B).

Comparison of the cryoEM maps for the apo and SAL-complexed HSA clearly indicated the presence of residual density in domain IB (also called FA1), IIA (FA7, or Sudlow site 1) and IIIa (FA3-FA4 cleft, or Sudlow site 2), that cannot be attributed to protein residues (Figure 4). Three SAL molecules were modeled in the three cryoEM sites (Figure 4). The assigned orientation is the most consistent with the available electron density, but given the relatively low resolution, alternative orientations are plausible. In the case of a novel binder, even approximate structural information may provide a starting point for ligand binding site identification and subsequent computational modeling and in silico optimization.

**Figure 4:**
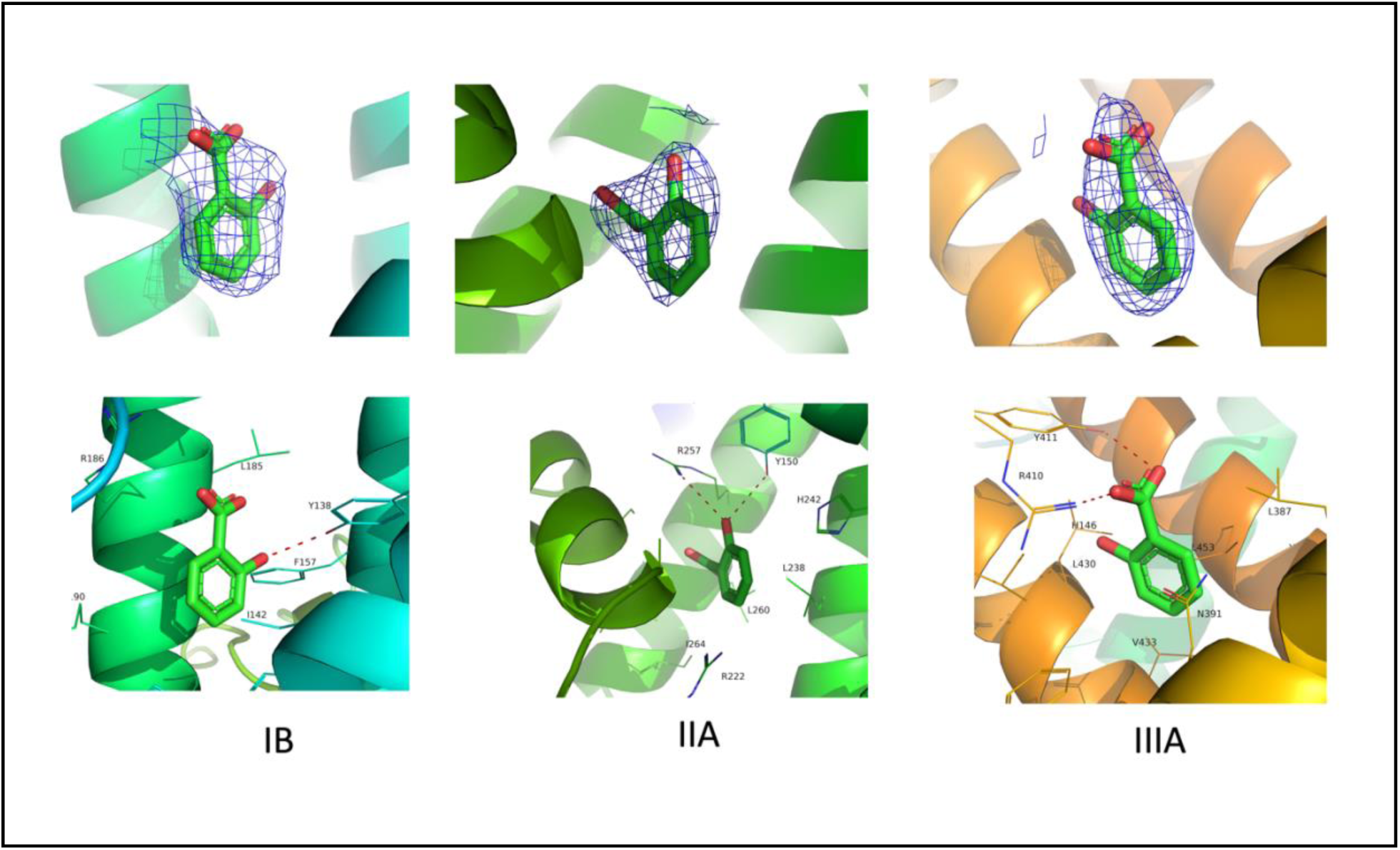
Salicylic acid binding sites in the cryoEM structure. Top row: residual density observed in the three sites, that was interpreted as bound SAL molecules. Bottom row: interactions made by the three SAL molecules in the three binding sites

While two of the SAL binding sites (in domain IB and IIA, Sudlow site 1) were previously identified in the crystal structures, the third binding site (in site IIIA, Sudlow site 2) is unique to the cryoEM structure. Sudlow site 1 has been demonstrated to prefer large heterocyclic and negatively charged compounds, whereas Sudlow site 2 is the preferred site for small aromatic carboxylic acids^42,46^, which is consistent with the cryoEM structure and other reported structures^4^. Interestingly, this pocket is occupied by small molecule binders only when the HSA is defatted. Subdomain IIIA has been reported to be a strong binding site for fatty acids^7^; it is likely that fatty acids pre-occupying this site may inhibit other drugs from competing efficiently and binding in this site. In the crystal structures, this site is indeed occupied by two myristate molecules (Figure 3B), a result of the procedure used to generate the samples and the crystals (soaking of crystals previously grown in presence of myristate^9^).

The Sudlow Site 1 (in domain IIA) is occupied by a SAL molecule in all structures. The orientation of the SAL molecule is approximately the same, although there is a slight shift in the position, related to the different conformation of the protein associated with the presence (in the crystallographic structures) of bound myristate and other ligands (Figure 3B). The site in domain 1B (FA1) is occupied in crystal structures 2I30 and 3B9M, and in the cryoEM structure. No SAL has been modeled in that site in structure 2I2Z, although bound myristate is present. In structure 2I2Z the salicylic acid is the result of the HSA driven hydrolysis of acetylsalicylic acid^9^, and the resulting concentration may be too low to compete with myristate in the FA1. The differences observed between the crystal structures and the cryoEM structures show that cryoEM allows us to investigate multiple binding sites without the conformational changes or competition effects of fatty acids, or the constraints induced by crystallization, thus providing different insights in HSA binding properties.

### CryoEM structure of teniposide bound to HSA

The final map for HSA in complex with teniposide was reconstructed to a nominal resolution of 3.4 Å (Figure 1, Table 1 and Table 2). There is one crystal structure of teniposide bound to HSA^3^ (PDB ID: 4L9Q); these crystals were grown in presence of teniposide but in the absence of fatty acids. Consequently, the overall conformation of teniposide-bound HSA closely resembles that of the apo protein (Figure 5). The most significant difference lies in the position of domain IIIB, which, based on the local resolution map, exhibits a high degree of flexibility. In the crystal structure, this domain is involved in extensive interactions between the two molecules identified in the asymmetric unit, likely limiting its mobility.

**Figure 5:**
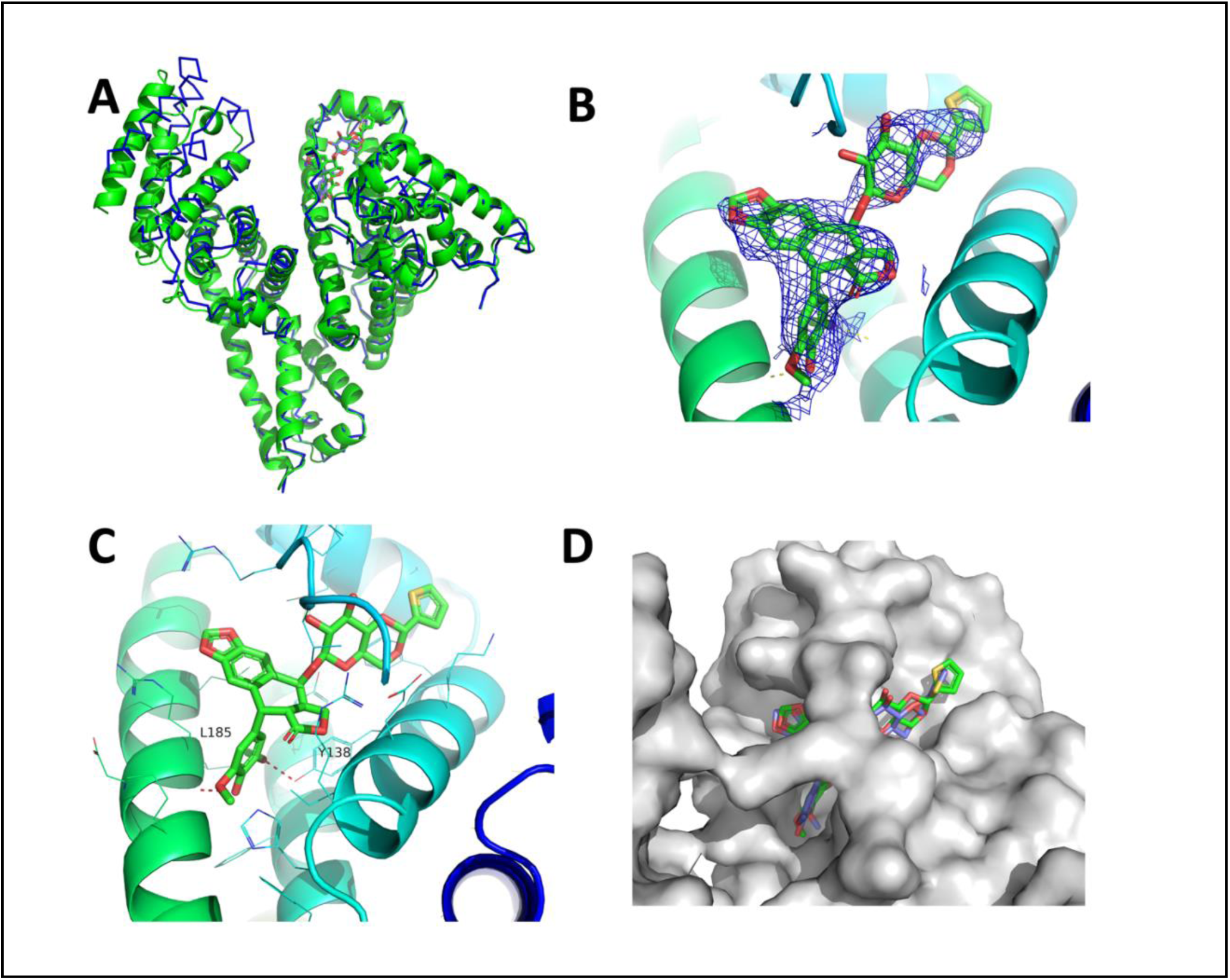
CryoEM structure of teniposide bound to HSA. **a** Overlay of the X-ray structure (PDB ID: 4L9Q, blue ribbon) and the cryoEM structure (green cartoon); **b** residual density observed in the domain 1B and attributed to teniposide; **c** teniposide binding site. **d** Overlay of the teniposide molecules from 4L9Q and the cryoEM structure. HSA (from the cryoEM structure), is shown as a gray surface

Like the X-ray structure, in the cryoEM structure, teniposide is situated in the highly hydrophobic binding pocket located in subdomain IB. As anticipated from the teniposide logP value (1.7), the binding is primarily driven by hydrophobic interactions and likely the displacement of water molecules (Figure 5C) that at the current resolution cannot be modeled. Interestingly, a couple of favorable polar interactions are facilitated by the 4-hydroxy-3,5-dimethoxyphenyl moiety of teniposide, specifically two hydrogen bond interactions: one between a methoxy-group of teniposide with the hydroxyl group of the phenol ring of tyrosine 138 (Y138) and the second one between the hydroxyl group of teniposide and the backbone of leucine 185 (L185) (Figure 5C).

Comparing the cryoEM structure to 4L9Q, it is clear that in both structures the central four-ring system (tetrahydro-5H-[2]benzofuro[6,5-f][1,3]benzodioxol-8-one) is fairly well ordered, and aligns quite well. The rest of the molecules (thenthenylidene-beta-D-glucopyranoside and the 4-hydroxy-3,5-dimethoxyphenyl moieties) show a rather high degree of diversity in the two structures, probably due to the fact that both substructures sit in solvent exposed areas (Figure 5D).

## Conclusions

While cryoEM structures of molecules smaller that 100 kDa in size are available^25–28,31^ they have been often the results of a one-off process that tested the current technology and software, or obtained by extensive manipulation of the sample itself^26,47–49^ but did not address the need for reproducibility in structure determination, as well as the necessity of identifying ligands bound, two essential requirement for the use of cryoEM derived information in drug discovery and development. In this paper we have shown that routine structural determination of molecules less than 70 kDa in size is possible, and that visualization of bound ligands is achievable. The work was carried out on a Glacios microscope, suggesting that 200kV instruments can be enough to provide structural support in drug discovery. In addition, the cryoEM workflow allows for the analysis of the sample in conditions not limited by crystal packing or methodology used (for example, soaking of compounds in a pre-loaded system). Protein flexibility can be easily visualized directly from the map and can be used to explain the protein biological activity without the need of a model. Exploration of different states (presence or absence of lipids, different buffers, different salt concentrations, different pH) and their effect on binding of drugs and other small molecules can also be carried out by cryoEM, without being limited by specific crystallization conditions. This may help the understanding of the complex binding characteristics of very important carriers such as HSA. Although the resolution of a cryoEM map may limit the accurate identification of atom positions, availability of structural data can greatly support in silico modeling to derive pharmacophore specific characteristics that enhance or limit bind of drug molecules to specific targets.

## Methods

### Sample preparation and cryo-electron microscopy

Human serum albumin (Sigma-Aldrich, Lot No.: SLBZ2785) was reconstituted in 1 ml of PBS to achieve a final concentration of 120 mg/mL. For the drug-bound samples, solutions of teniposide (Enzo, Lot No.: 10212202) in DMSO and salicylic acid (Thermo Scientific, Lot No.: 102430) in EtOH were added, resulting in a final concentration of 50 mM, while ensuring that the final DMSO/EtOH concentration did not exceed 1%. The mixtures were then incubated overnight at 4℃. Subsequently, the three samples were diluted to a concentration of 6.0 mg/mL using their respective stock solutions. A volume of 3.0 µL of each sample was applied to glow discharged (PELCO easiGlow™) 300 mesh holey gold UltrAuFoil R 1.2/1.3 grids. Grid preparation was performed using a Vitrobot Mark IV (Thermo Fisher Scientific, Eindhoven, The Netherlands), with the environmental chamber set to 95% humidity and a temperature of 4℃. The specific blotting conditions are detailed in Table 1.

Imaging was conducted using a Glacios Cryo-Transmission Electron Microscope (Thermo Fisher Scientific, Eindhoven, The Netherlands) equipped with a Falcon 4 camera (Thermo Fisher Scientific, Eindhoven, The Netherlands). Movies were recorded using the Leginon software version 3.6^50^ at a nominal magnification of 240,000x with a calibrated pixel size of 0.575 Å for the unbound HSA (0.566 Å for the drug-bound samples) and a dose rate of 10.41 e^-^/Å^2^/s with a total exposure of 3.50 seconds, for an accumulated dose of 36.43 e^-^/Å^2^. Movies were collected at a nominal defocus range of −0.5 µm to −1.5 µm.

### CryoEM image processing

For all datasets, data processing procedures followed highly similar strategies. The apo HSA will be used as an example of the general workflow. On-the-fly data preprocessing and quality control was performed within cryoSPARC live^51^. Patch motion correction^51^ was used for correcting beam-induced motion and to account for stage drift. The contrast transfer function was estimated for each micrograph using patch CTF^52,53^. Micrographs with contrast transfer function fits lower than 5 Å were removed. 418,782 particles were picked with a Topaz pre-trained neural network model^54^ and extracted with a box size of 440 pixels from 3,271 manually curated micrographs. All subsequent steps were carried out in Cryosparc^51^. An initial clean-up of the extracted particles was performed by iterative rounds of 2D classifications, yielding 105,192 particles. The selected particles were used in an ab-initio reconstruction followed by heterogenous refinement. The best resolved model after heterogeneous refinement, 54,988 particles, was used for a non-uniform (NU) refinement as implemented in Cryosparc^51^. This map was selected for further refinement, carried out using CTF local and global refinement followed by local resolution refinement. The estimated resolution is 3.5 Å based on the gold standard Fourier shell correlation of 0.143. For a comprehensive overview of the processing workflow, please refer to Figure 2 and Table 2.

### Model Building and refinement

Model building and refinement were initiated with a published model for HSA (PDB ID:1AO6). The initial structure was placed into the sharpened density maps using the ChimeraX “fit to map” utility^55^. Generation of cartesian coordinates and geometry restraints for salicylic acid and teniposide were carried out in eLBOW from their respective SMILE strings^56^. Iterative rounds of model building and refinement were performed in PHENIX v.1.19.2^57^ and COOT v. 0.9.6 EL^58^. The final models were validated against the half-maps and its quality assessed by MolProbity^59^. For a full summary of the refinement statistics see table 2.

## Data availability

EM maps and atomic models were deposited to the Electron Microscopy Data Bank (EMDB) and Protein Data Bank (PDB) databases. PDB codes for the various structures reported in this manuscript are 8VAF, 8VAE, 8VAC and the EMDB accession codes are EMD-43090, EMD-43089, EMD-43088 for HSA apo, in complex with salicylic acid, and in complex with teniposide respectively.

## Author contributions

K.W.L., S.S., N.L.T., A.C.F. and S.M.S. prepared the grids. K.W.L., S.S., N.L.T., A.C.F. collected the data. EM processing work was carried out by C.C. and C.C. built and refined the structure. D.T. and P.V.D. were involved in grid making and data collection in the initial phase of the project. C.C., N.P. and G.S. supervised the work and wrote the manuscript. All authors read the manuscript and discussed the materials.

## Notes

### Competing Interest Statement

The authors have declared no competing interest.

