## Supplementary information for "The CryoEM structure of human serum albumin in complex with ligands"

Establishment of general vitrification conditions. After reconstituting the purchased HSA in phosphate-buffer saline pH 7.4 (PBS), initial quality control assessment of the sample was extensively carried out; size exclusion chromatography, mass photometry, and negative staining were used to evaluate factors such as precipitation, aggregation, and stability of the sample. (Supp Figure 1). The sample's performance after extended storage at both 4°C and -80°C under various concentrations was also examined. The results clearly indicated the presence of a monomeric, highly homogeneous, and monodisperse sample under a variety of conditions.

As a subsequent step, our focus shifted towards implementing a robust 3D structure determination workflow, encompassing the entire process from vitrification to structure solution for the unliganded sample, to exploring a diverse range of conditions, including variations in protein concentration, grid type, and vitrification methods (as detailed in Supp Table 1). The objective was to pinpoint the most effective concentration, grid type, and vitrification parameters, while simultaneously evaluating the reproducibility of the method. These insights could then serve as a foundational reference for vitrifying complexes in subsequent experiments.

All grids underwent screening on a Thermo Fisher Scientific Glacios Cryo-Transmission Electron Microscope (cryo-TEM) equipped with a Falcon 4 direct electron detector, operating at 200 kV. The initial assessment included evaluations of ice quality, particle distribution, and particle behavior under varying ice thickness. For grids meeting suitability criteria, high-resolution data collections were carried out. In total, 83 grids were screened, resulting in the collection of 14 data sets, allowing for the evaluation of different data collection procedures. CryoSPARC[1] was employed for the processing of all data sets, aiding in the identification of potential issues such as preferred orientation, aggregation, and protein deterioration at the water/air interface. This iterative cycle of vitrification, screening, data collection and data processing was repeated multiple times until a set of vitrification conditions, data collection parameters (as outlined in Table 1), and data processing steps (depicted in Figure 2) were identified, ultimately yielding a 3.5 Å map for the unliganded protein.

### Supplementary tables

| Sample Name | Concentration (mg/ml) | Grid Type | Hole size (microns) | Vitrification |
|---|---|---|---|---|
| HSA (SD) - Diluted with PBS | 3.604 | UltrAuFoil | 1.2x1.3 | V: Vitrobot #2 bt/bf/wt=8.0/2/20 100%/4C Solarus 10.0s |
| HAS 0.05% Glutaraldehyde - SEC purified dimer, concentrated fractions | 6.000 | UltrAuFoil | 1.2x1.3 | V: Vitrobot #4 bt/bf/wt=10.0/10/0 95%/4C EasiGlow2 60.0s |
| HAS 0.05% Glutaraldehyde - SEC purified dimer, unconcentrated fraction D8 | 3.200 | UltrAuFoil | 1.2x1.3 | V: Vitrobot #4 bt/bf/wt=10.0/10/0 95%/4C EasiGlow2 60.0s |
| HSA + TCEP 100mM 1.27 mM + BMH 20mM 0.30 mM | 9.641 | Nanowire-C | | C: Chameleon: 190.0ms plasma cleaning 60.00s |
| HSA + TCEP 100mM 1.27 mM + BMH 20mM 0.30 mM | 3.856 | UltrAuFoil | 1.2x1.3 | V: Vitrobot #3 bt/bf/wt=6.0/0/0 95%/4C EasiGlow2 60.0s |
| HSA + TCEP 100mM 1.27 mM + BMH 20mM 0.30 mM | 3.856 | C-Flat Cu-Mesh | 2x1 | V: Vitrobot #3 bt/bf/wt=10.0/-5/0 95%/4C EasiGlow2 60.0s |
| HSA + TCEP 100mM 1.27 mM + BMH 20mM 0.30 mM | 3.856 | C-Flat Cu-Mesh | 1.2x1.3 | V: Vitrobot #3 bt/bf/wt=10.0/-5/0 95%/4C EasiGlow2 60.0s |
| HSA | 34.000 | Nanowire-C | | C: Chameleon: 200ms plasma cleaning 60.00s |
| HSA | 1.000 | C-Flat Cu-Mesh | 1.2x1.3 | V: Vitrobot #3 bt/bf/wt=16.0/13/20 100%/4C EasiGlow2 10.0s |
| HSA | 0.010 | Continuous carbon grid (NIS) | | N: UF4 drop/3 uL - Rinse: 5 drop/3 uL EasiGlow2 20.0s |
| HSA | 2.001 | C-Flat Cu-Mesh | 1.2x1.3 | V: Vitrobot #3 bt/bf/wt=16.0/13/20 100%/4C EasiGlow2 10.0s |
| HSA | 2.001 | UltrAuFoil | 1.2x1.3 | V: Vitrobot #3 bt/bf/wt=16.0/0/20 100%/4C EasiGlow2 20.0s |
| HSA | 9.999 | Nanowire-C | | V: WBG Chameleon 145 ms plasma cleaning 60.00s |
| HSA | 3.000 | C-Flat Cu-Mesh | 1.2x1.3 | V: Vitrobot #3 bt/bf/wt=6.0/0/0 95%/4C EasiGlow2 60.0s |
| HSA | 2.000 | C-Flat Cu-Mesh | 1.2x1.3 | V: Vitrobot #3 bt/bf/wt=6.0/0/0 95%/4C EasiGlow2 60.0s |
| HSA | 4.001 | C-Flat Cu-Mesh | 1.2x1.3 | V: Vitrobot #3 bt/bf/wt=10.0/-5/0 95%/4C EasiGlow2 60.0s |
| HSA | 4.001 | C-Flat Cu-Mesh | 1.2x1.3 | V: Vitrobot #3 bt/bf/wt=6.0/0/0 95%/4C EasiGlow2 60.0s |
| HSA | 4.000 | C-Flat Cu-Mesh | 1.2x1.3 | V: Vitrobot #4 bt/bf/wt=3.0/10/0 95%/4C EasiGlow2 60.0s |
| HSA | 4.000 mg/mL | C-Flat Cu-Mesh | 1.2x1.3 | V: Vitrobot #4 bt/bf/wt=8.0/10/0 95%/4C EasiGlow2 60.0s |
| HSA | 4.000 mg/mL | UltrAuFoil | 1.2x1.3 | V: Vitrobot #4 bt/bf/wt=8.0/10/0 95%/4C EasiGlow2 60.0s |
| HSA | 8.000 mg/mL | UltrAuFoil | 1.2x1.3 | V: Vitrobot #4 bt/bf/wt=10.0/10/0 95%/4C EasiGlow2 60.0s |
| HSA | 4.000 mg/mL | C-Flat Cu-Mesh | 1.2x1.3 | V: Vitrobot #4 bt/bf/wt=10.0/10/0 95%/4C EasiGlow2 60.0s |
| HSA | 4.000 mg/mL | C-Flat Cu-Mesh | 1.2x1.3 | V: Vitrobot #4 bt/bf/wt=10.0/10/0 95%/4C EasiGlow2 60.0s |
| HSA | 0.020 | Continuous carbon grid (Ted Pella) | | N: UF4 drop/3 uL - Rinse: 5 drop/3 uL EasiGlow2 20.0s |
| HSA | 0.010 | Continuous carbon grid (Ted Pella) | | N: UF4 drop/3 uL - Rinse: 5 drop/3 uL EasiGlow2 20.0s |

### Supplementary Figures

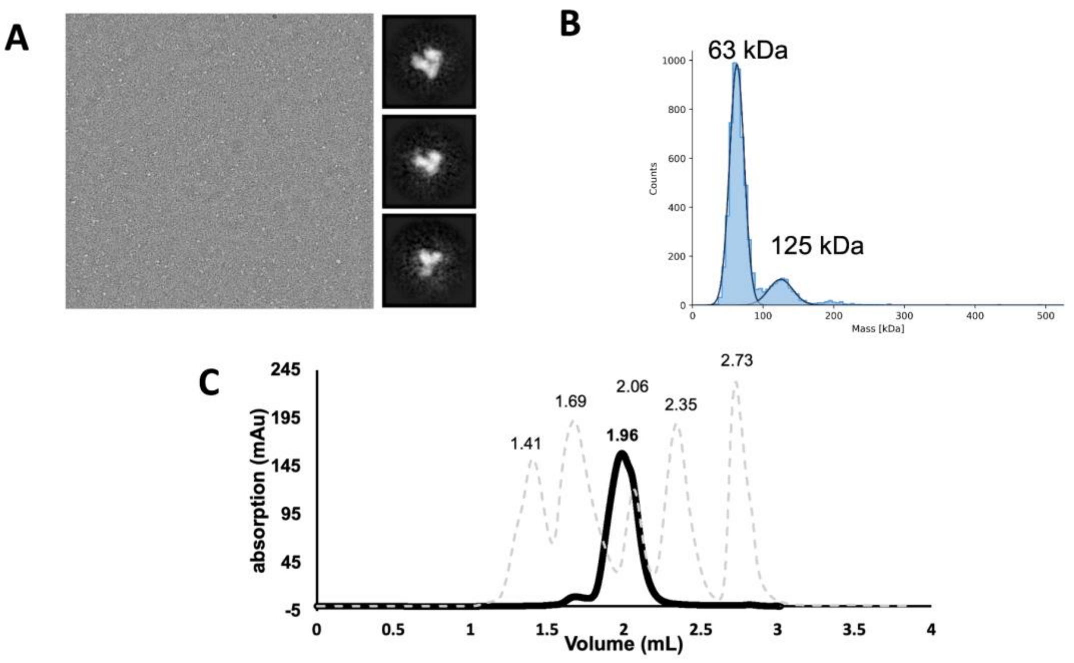


**Supplementary Fig. 1. Quality control results for HSA reconstituted in PBS pH=7.4 a** Negative stain micrograph for HSA with corresponding 2D classes average. **b** Mass distributions as measured using mass photometry (Refeyn Ltd.).**c** Size exclusion chromatogram showing the comparison between sample (solid black line) and calibration (broken grey line). Calibration peaks: # 1.41 Thyroglobulin (bovine) 670 kDa; # 1.69 $\gamma$-globulin (bovine) 158 kDa; # 2.06 Ovalbumin (chicken) 44 kDa; # 2.35 Myoglobin (horse) 17 kDa; # 2.73 Vitamin B12 1.35 kDa

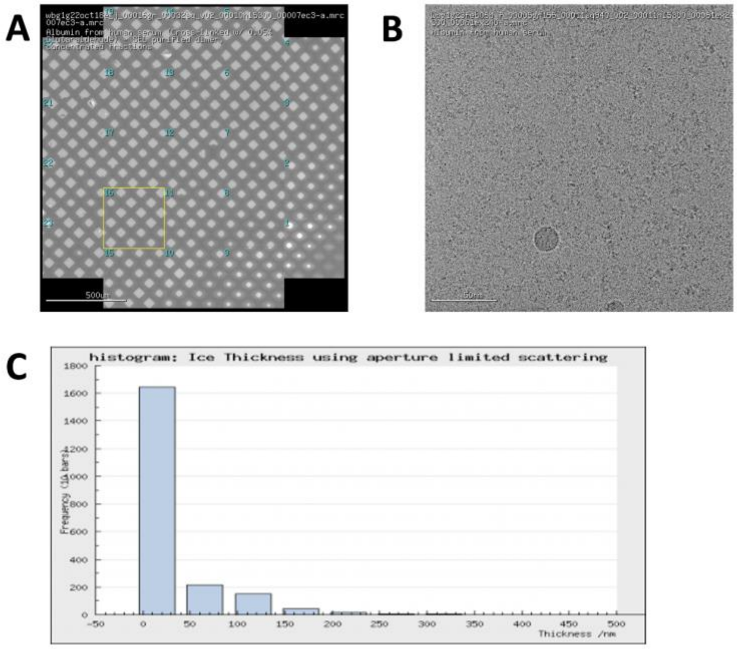

**Supplementary Fig. 2. Grid quality and ice thickness for the grid that yielded the 3.5 Å structure of apo HSA. a** overview of the low magnification atlas. **b** Exemplary micrographs. **c** Plot of ice thickness as evaluated in Appion[2]. **a** and **b** are from Leginon software [3,4].

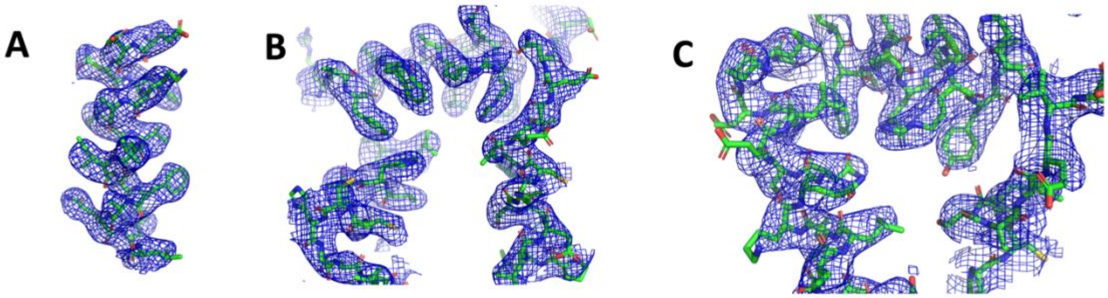


**Supplementary Fig. 3. An example of the quality of the map for the apo HSA.** A) helix spanning residues 16-31; B) the Sudlow site 1 C) Sudlow Site 2. Figure generated with Pymol[5]

5. Schrödinger, L. The PyMOL Molecular Graphics System, Version~1.8.
